# Using Shape Fluctuations to Probe the Mechanics of Stress Granules

**DOI:** 10.1101/2022.05.03.490456

**Authors:** Jack O. Law, Carl M. Jones, Thomas Stevenson, Matthew S. Turner, Halim Kusumaatmaja, Sushma N. Grellscheid

## Abstract

Surface tension plays a significant role in many functions of biomolecular condensates, from governing the dynamics of droplet coalescence to determining how condensates interact with and deform lipid membranes and biological filaments. To date, however, there is a lack of accurate methods to measure the surface tension of condensates in living cells. Here, we present a high-throughput flicker spectroscopy technique that is able to analyse the thermal fluctuations of the surfaces of tens of thousands of condensates to extract the distribution of surface tensions. Demonstrating this approach on stress granules, we show for the first time that the measured fluctuation spectra cannot be explained by surface tension alone. It is necessary to include an additional energy contribution, which we attribute to an elastic bending rigidity and suggests the presence of structure at the granule-cytoplasm interface. Our data also show that stress granules do not have a spherical base-shape, but fluctuate around a more irregular geometry. Taken together, these results demonstrate quantitatively that the mechanics of stress granules clearly deviate from those expected for simple liquid droplets.

## Introduction

Biomolecular condensates are cellular compartments which lack a bounding membrane. They form through a process of liquid-liquid phase separation (1), and are increasingly associated with a wide range of biological functions. Importantly, their ability to carry out these functions is closely related to their mechanical properties. For example, condensates are thought to act as reaction crucibles and sites of chemical sequestration (1). This has led to an interest in studying their diffusion constants and viscosities, as these determine the rates of biochemical reactions. In addition, dysregulation of condensates leading to a liquid-to-solid phase transition due to the formation of fibrils has been implicated in numerous neurodegenerative diseases such as ALS (amyotrophic lateral sclerosis) (2). More recently, there is growing recognition of the mechanical significance of the condensate surface tension, as condensates exert forces on each other, on lipid membranes and on cytoskeletal filaments (3, 4).

Characterising the relevant mechanical properties of condensates in living cells is highly challenging. In particular, accurate techniques to measure surface tension are currently lacking. Standard surface tension methods such as sessile drop tensiometry, Du Noüy ring or Wilhelmy plate (5), while highly accurate for measurements in-vitro, are not applicable in living cells. To date, the surface tension of condensates in living cells has been estimated from the timescale of droplet coalescence events (6). Typically only a handful of events are analysed because identifying them in live cell-cell imaging is cumbersome. Furthermore, this approach employs a gross assumption that such a timescale depends on a simple viscosity-surface tension ratio (7) and it requires the condensate viscosity to be known. Measurements of condensate viscosity are often based on fluorescence recovery after photobleaching (FRAP), which itself is known to have a number of drawbacks, including a strong dependence on the area bleached and choice of fluorescence recovery model (8). Hence, the droplet coalescence approach can only provide an order of magnitude at best. It is not precise or sensitive enough to track the evolution or variation in condensate surface tension in living cells. In addition, it cannot distinguish between surface tension and other mechanical forces influencing the coalescence dynamics. There is a clear need for novel approaches for accurate and direct measurements of surface tension.

We present flicker spectroscopy as a powerful solution to this problem. The basis of flicker spectroscopy is the analysis of the thermal fluctuations of the condensate shapes. In contrast to previous studies (7), here we undertake detailed spectral analysis of the fluctuations using stress granules as a model condensate organelle. The presented approach, however, should be widely applicable to other condensates in living cells. Harnessing flicker spectroscopy, we reveal that the mechanics of stress granules depend on a previously-unreported elastic bending deformation, and their base-shape is generally non-spherical.

## Results and Discussion

Stress granules were induced using 200μM sodium arsenite in U2OS cells stably expressing G3BP1-GFP and lacking endogenous G3BP1 and G3BP2 (9) (Fig. 1A-B). They were then imaged using a spinning-disk microscope for forty seconds, and a bespoke edge detection algorithm was developed and used to detect the boundary of each granule on the imaging plane, as illustrated for an example granule in Fig. 1C. The collected granule boundaries across the entire measurement period, sampled every 10 ms, is shown in Fig. 1D. The shape deformation in each granule boundary was then decomposed into its independent Fourier modes *q*, as shown in Fig. 1G, and we time-average the squared amplitude of each mode 〈|υ_*q*_|^2^〉. Importantly, the spectra observed from live cells are distinct from those from identical fixed cells, demonstrating that the observed fluctuations are not just due to microscope noise, as illustrated in Fig. 1E, for *q* = 2. Additionally, we also simulated randomly rotating rigid bodies with quenched shape roughness. As shown in Fig. 1F, this has a different spectrum and fits poorly when compared to the model discussed below, suggesting that the spectra measured from live cells are meaningful. A detailed analysis protocol is provided in Supplementary Material section 6 (SM. 6).

**Figure 1.**
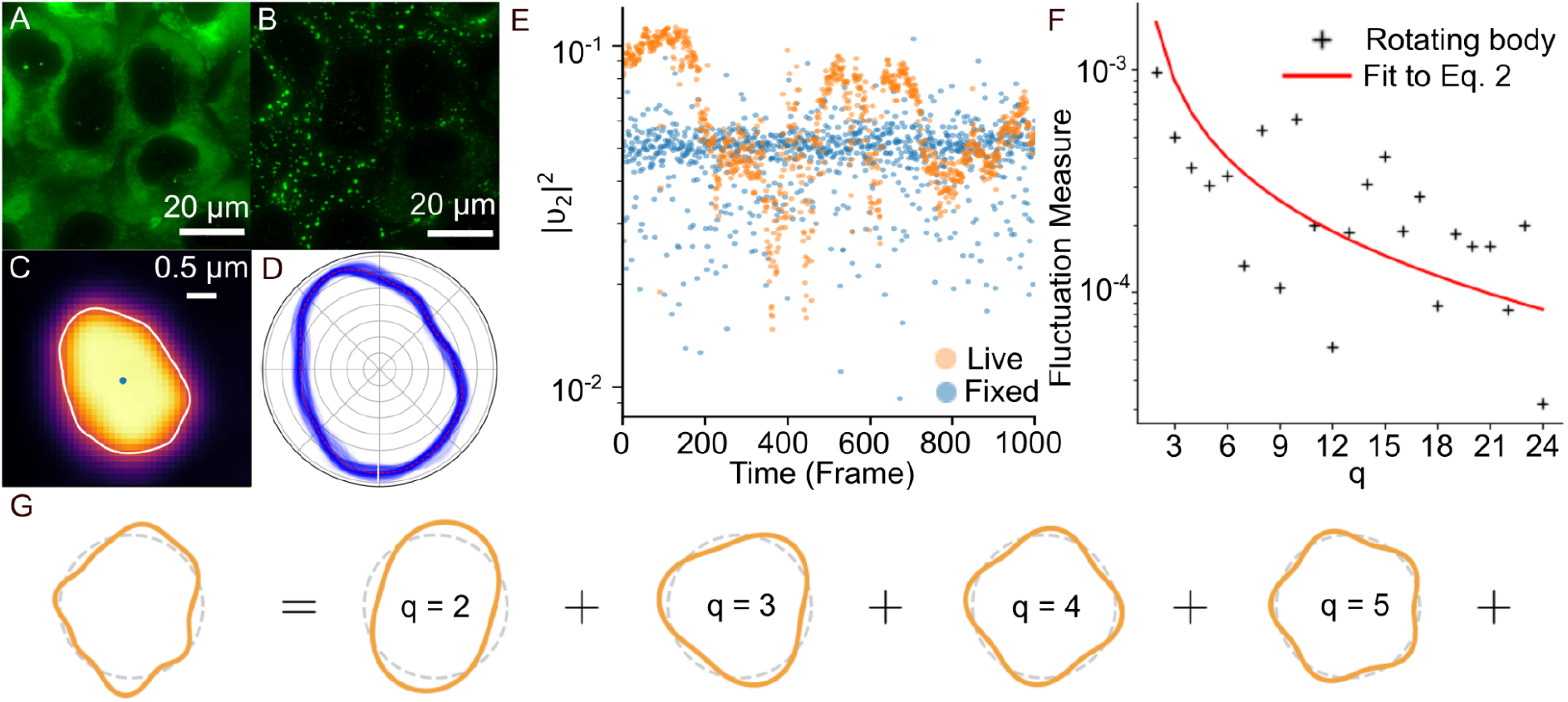
Analysing the shape fluctuations of stress granules for live cell flicker spectroscopy. (A) Unstressed U2OS cells, with stress granule component G3BP fluorescing green. (B) The same cells after arsenite treatment. G3BP has localised into stress granules. (C) A single stress granules is located and outlined by the boundary-tracking routine. (D) The fluctuations of the same stress granule are shown for each frame of the analysis. (E) In a live cell, the condensate boundary shows correlated fluctuations over time (orange), whereas in a fixed cell, only microscope noise is seen (blue). (F) In a simulated, non-fluctuating granule undergoing random rotations, the model does not fit to the measured spectrum. (G) A schematic showing the decomposition of a granule boundary into Fourier modes.

To infer the condensate surface tension and bending rigidity, the measured fluctuation spectra must be fitted to a theoretical prediction. Here, we find they fit well to a spectrum derived from the Helfrich Hamiltonian (10)

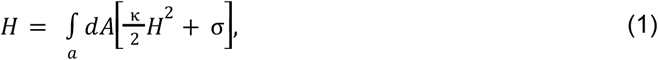

where *κ* is the bending rigidity, *H* is the total curvature and *σ* is the surface tension. This formulation includes the assumption that the spontaneous curvature of the surface is 0. For a simple liquid droplet, we would expect no contribution from bending rigidity. However, importantly, a number of works have recently suggested the presence of structure at the condensate interface, such as intrinsically disordered proteins acting as Pickering stabilisers on P-granules (11) and FXR1 proteins being enriched on stress granules (12). When projected to the two-dimensional imaging plane, as is the case in our setup, the time-average fluctuation spectra are given by (13)

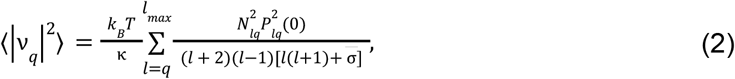

where υ_*q*_ is the magnitude of the *q*^th^ fluctuation mode, *k_B_* is the Boltzmann constant, *T* is the temperature, *P_lq_* and *N_lq_* are the associated Legendre polynomials and normalisation factor respectively and 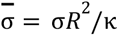 is the dimensionless surface tension, with *R* the condensate radius (see SM. 1 for a detailed derivation). *l_max_* = 75 is sufficient for the sum to converge. An example of such a fit can be seen in Fig. 2A.

**Figure 2.**
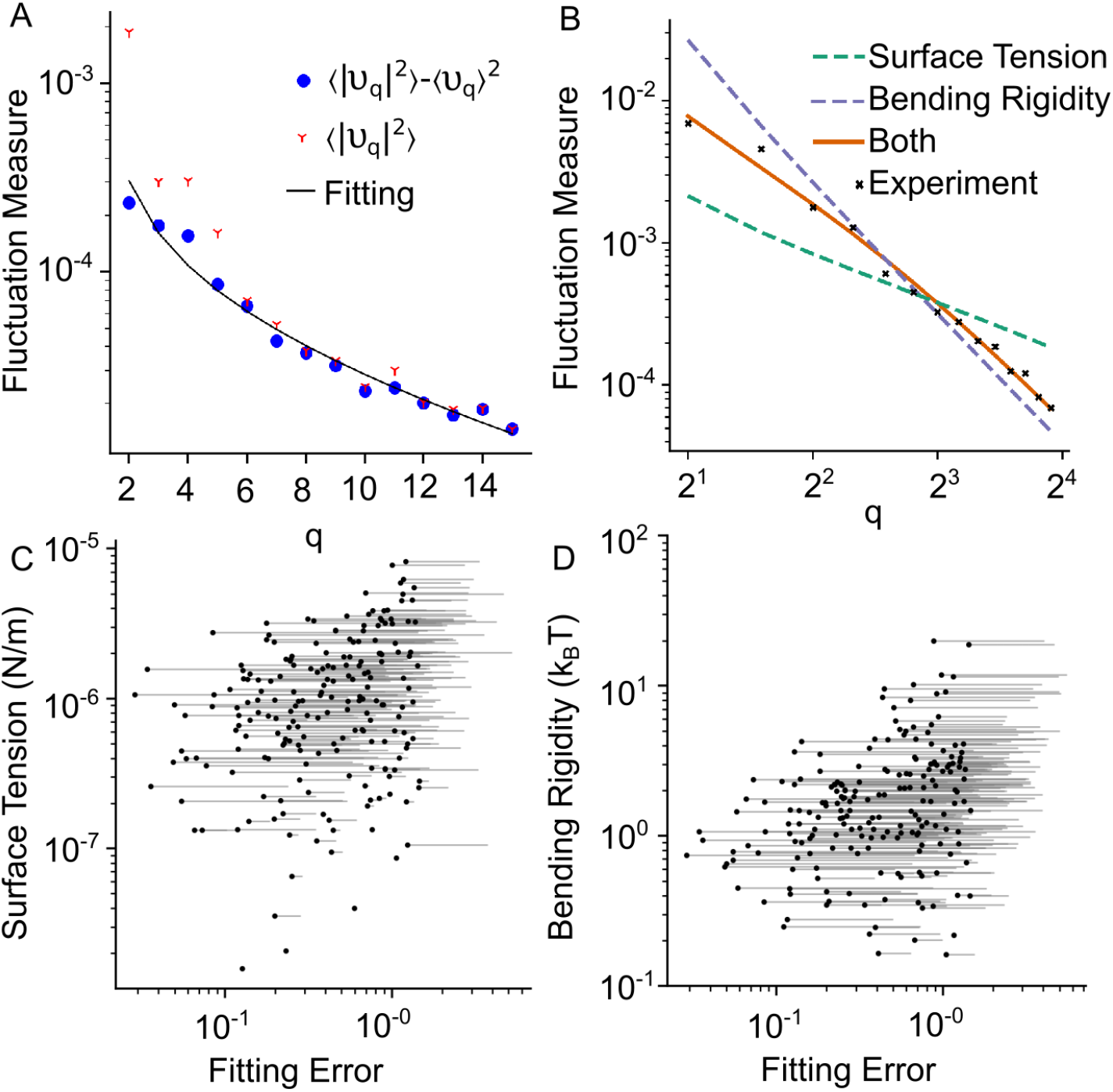
Using flicker spectroscopy to extract the mechanical properties of stress granules. (A) The measured fluctuation spectrum of an example stress granule (red stars), the spectrum after correction for the non-spherical base shape (blue dots) and the fitted model spectrum (black line). (B) Another example fluctuation spectrum (black crosses) with fitting to three theoretical spectra: surface tension only (green dashes), bending rigidity only (purple dash) and both surface tension and bending rigidity (orange line). Only the model with both surface tension and bending rigidity fits well to the experimental data. (C) The change in fitting error (solid line) when the bending rigidity is removed. For clarity, we select 300 granules at random. For each point the fitting is worse without bending rigidity. (D) Similar to (C), the change in fitting error when the surface tension is removed. Again, the fitting is worse.

We collected microscopy videos of 99091 granules over several independent experimental runs. After fitting, we filtered out all granules lacking a closed outline in at least 60% of the frames. These were granules typically undergoing merging, or they may have wetted around a sub-cellular surface. These 49415 granules were further filtered out if the error on the fit to the theoretical spectrum was greater than 0.6 (see SM. 5), resulting in a final selection of 14973 stress granules.

A comparison of the measured and predicted spectra of the stress granules above led to the following insights. Firstly, if stress granules are simple liquid droplets, their fluctuations are expected to depend mainly on surface tension. However, as shown in Fig. 2B, we observed that the resulting fit from including surface tension alone (setting *κ* = 0) was very poor. This is especially so for higher order *q* modes, where the bending term was expected to dominate from Eq. 2. Similarly, the measured and predicted spectra are in poor agreement if only elastic bending deformation is taken into account (setting σ = *0*), especially for the lower *q* modes. Fig. 2C and 2D quantify the increase in fitting errors when either surface tension or bending rigidity is ignored.

Secondly, we find that the base shape of the stress granules is not perfectly spherical, at least over the duration of the measurement, further discrediting a simple-liquid model of stress granules. This issue can be detected by showing that the average 〈υ_*q*_〉 is non-zero. The fitting routine must be amended to account for this, which can be achieved by subtracting the time-independent part of the spectrum (13), meaning that we instead analyse the following fluctuation measure

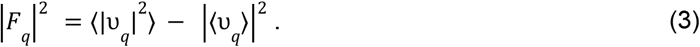

Failing to account for this base-shape correction leads to a spectrum that fits poorly to the theoretical model (Fig. 2A), and to significant misestimation of the surface tension and bending rigidity (Fig. 3A-B), including falsely identifying many droplets with negligible surface tensions. Using the full, corrected, model, we find the measured surface tension spans across several orders of magnitude, primarily between the orders 0.1 μN/m and 10 μN/m, with an average of (0.62±0.02) μN/m. The obtained range of values is in fact in excellent agreement to previous estimates in the literature for the nucleolus and germline P-granules (3, 7, 14). The bending rigidity was found to be (1.9±0.05) *k_B_T*, which is consistent with thermodynamically stable aggregates at the interface (15).

**Figure 3.**
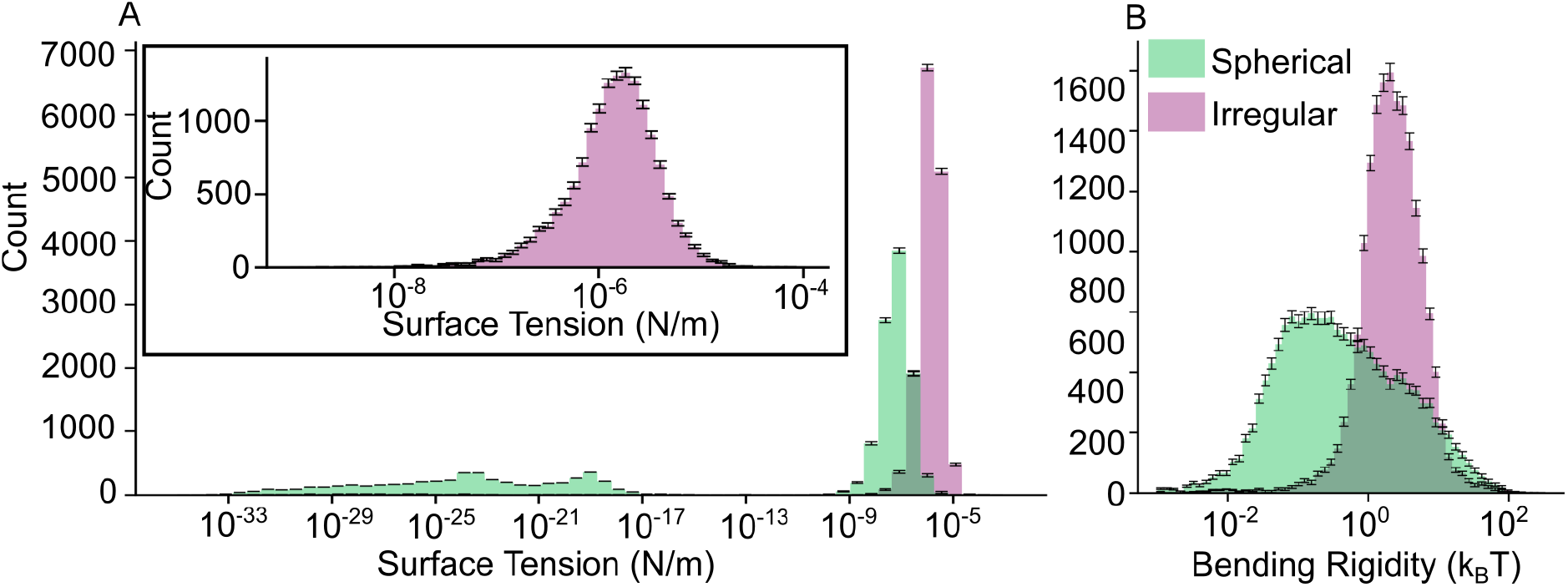
The distribution of surface tension and bending rigidity in stress granules. Histograms of the surface tension (A) and bending rigidity (B), with (purple) and without (green) the correction for non-spherical base shape given in Eq. 3. Included in the histograms are all granules with a fitting error higher than 0.6 and a success rate greater than 0.6 from the analysis of the corrected spectra, so all histograms contain the same 14973 granules. Not including the correction causes a severe underestimate of the surface tension, and results in many granules being falsely attributed a negligible surface tension. The inset in (A) shows a more detailed view of the surface tension histogram with the shape-correction.

In conclusion, we have demonstrated live-cell flicker spectroscopy as a high-throughput approach for exploring two key mechanical properties of condensate organelles under physiological conditions. We demonstrate here for the first time that bending rigidity is an essential, measurable property of stress granules and demonstrate the importance of accounting for the non-spherical shape when calculating condensate mechanical properties. We believe our work opens the way for a number of key advantages of live cell flicker spectroscopy to be harnessed, for example to evaluate the changes in mechanical properties of condensates in living cells as a function of time, in the presence of disease relevant mutations, or in response to candidate drugs. At the same time, there remains a number of exciting avenues to extend our analysis, including to infer the condensate viscosity/viscoelasticity from the temporal variation of fluctuation modes, and to consider the possible effects of bulk elasticity and active mechanical forces.

## Materials and Methods

All data, materials and equations needed to evaluate the conclusions in the paper are provided in the paper. Additional data related to this manuscript may be requested from the authors.

## Acknowledgments

We thank Nancy Kedersha, Harvard University (retired) for many insightful discussions. We thank Tim Hawkins, at the Durham University Microscopy unit and Hege Dale at the Molecular Imaging Centre, University of Bergen, Norway for helpful discussions. S.N.G, H.K and C.M.J acknowledge funding and support from the Biophysical Sciences Institute, Durham University. S.N.G acknowledges support from the Trond Mohn Stiftelse (No. BFS2017TMT01) and The Royal Society. H.K acknowledges support from EPSRC (Grant No. EP/V034154/1)

