## Supplementary Materials for "Using Shape Fluctuations to Probe the Mechanics of Stress Granules"

### 1. Derivation of the theoretical spectrum

In this section we discuss the derivation of the theoretical spectrum given in eq. 1 of the main text. The surface of a granule, $S$, is described as a perturbation, $u$, on the surface of a sphere radius $R$,

$$S(\theta, \varphi, t) = R[1 + u(\theta, \varphi, t)]. \tag{S1}$$

We may then write the perturbation as a weighted sum of spherical harmonics $Y_{lm}$,

$$u(\theta, \varphi, t) = \sum_{l=2}^{\infty} \sum_{m=-l}^{l} Y_{lm}(\theta, \varphi) U_{lm}(t), \tag{S2}$$

where $U_{lm}(t)$ is the magnitude of the given spherical harmonic term. $l$ and $m$ are the indices of the spherical harmonic modes. In general, $l \geq 0$ and $-l \leq m \leq l$. The $l = 0$ mode corresponds to a uniform (spherical) growth and shrinking. The time-average of this mode simply gives the average radius of the stress granule, which should be constant for a fixed-volume granule. It is represented by the first term in Eq. S1. The $l = 1$ mode represents translations of the whole granule, so has no bearing on thermal fluctuations of the surface. Therefore, we can discard these terms.

Assuming the energies to deform the condensate are as given by the Helfrich Hamiltonian, the average energy for each fluctuation mode is given by (1, 2)

$$\langle E_{lm} \rangle = \frac{\kappa}{2} \left| U_{lm} \right|^2 (l + 2)(l - 1)[l(l + 1) + \bar{\sigma}], \tag{S3}$$

where $\bar{\sigma} = \sigma R^2/\kappa$ is the dimensionless surface tension, with $R$ the mean condensate radius. The equipartition theorem states that all these modes should have the same time-averaged energy $\langle E_{lm} \rangle = k_B T/2$, giving an expected spectrum of

$$\langle \left| U_{lm} \right|^2 \rangle = \frac{k_B T}{\kappa} \frac{1}{(l + 2)(l-1)[l(l+1) + \bar{\sigma}]}. \tag{S4}$$

In practice, we are only able to measure a two-dimensional slice of the surface, so we must relate the three-dimensional fluctuation modes $\langle \left| U_{lm} \right|^2 \rangle$ into terms that we can observe directly. As we assume that the granules are imaged through the equatorial plane of the granule, then we can see a cross-section of the shape perturbations $\hat{u}(\varphi, t) = u(2\pi, \varphi, t)$ without loss of generality, giving the $\theta = \frac{\pi}{2}$ plane.

We can relate $\hat{u}$ to the two-dimensional fluctuation spectrum, by writing it as a sum of Fourier modes

$$\hat{u}(\varphi, t) = \sum_{q=2}^{\infty} \nu_q e^{-iq\varphi},$$

$$(S5)$$

where $\nu_q$ is the amplitude for mode $q$. Using standard Fourier transform, we can calculate $\nu_q$ via

$$\nu_q(t) = \frac{1}{2\pi}\int_0^{2\pi} \hat{u}(\varphi, t)e^{iq\varphi}d\varphi. \qquad (S6)$$

Importantly, this cross-section of the surface must match the definition of the full surface in eq. S2. This constraint leads to

$$\nu_q = \frac{1}{2\pi}\int_0^{2\pi} \sum_{lm} U_{lm}Y_{lm}(\pi/2, \varphi)e^{iq\varphi}d\varphi,$$

$$\nu_q = \frac{1}{2\pi}\int_0^{2\pi} \sum_{lm} U_{lm}N_{lm}P_{lm}\cos(\pi/2)e^{im\varphi}e^{iq\varphi}d\varphi, \qquad (S7)$$

where $P_{lm}$ and $N_{lm}$ are the associated Legendre polynomials and normalisation factor

$$N_{lm} = \sqrt{\frac{2l+1}{4\pi}\frac{(l-m)!}{(l+m)!}}, \qquad (S8)$$

respectively. Solving eq. S7 (2, 3), gives a relation between the time averaged magnitudes of the observable fluctuations $\nu_q$ and the full three-dimensional fluctuations $U_{lm}$

$$\langle|\nu_q|^2\rangle = \sum_{l=q}^{l_{max}} \langle|U_{lq}|^2\rangle N_{lq}^2P_{lq}^2(\cos\ \pi/2). \qquad (S9)$$

We take $l_{max} = 75$ to ensure a good convergence (2, 4). The result is usually accurate to within 0.1% after 40 terms. Substituting eq. S4 into eq. S9, we obtain

$$\langle|\nu_q|^2\rangle = \frac{k_BT}{\kappa}\sum_{l=q}^{l_{max}} \frac{N_{lq}^2P_{lq}^2(0)}{(l+2)(l-1)[l(l+1)+\bar{\sigma}]}, \qquad (S10)$$

which is the predicted spectrum of the experimentally observable fluctuations.

### 2. Cell culture

We used modified U2OS cell lines created by Nancy Kedersha called ΔΔ17-U2OS-EGFP-G3BP, ΔΔ17-U2OS- EGFP -G3BP2a and ΔΔ17-U2OS- EGFP -G3BP2b (5). These cells have endogenous G3BP1, G3BP2a and G3BP2b removed and the fluorescently labelled GFP-G3BP1, GFP-G3BP2a and GFP-G3BP2b added respectively. Cells were grown in T75 and T25 flasks containing Dulbeco’s Modified Eagle Medium (DMEM) (Sigma no. D5671). The cells were stressed in 35mm glass dishes (μ-Dish 35mm high Glass bottom, IBIDI Cat. No. 81158). For the arsenite treatment, 1 ml of 400 μM of sodium arsenite diluted in 1ml DMEM is added to 1 ml of untreated DMEM for a concentration of 200 μM. We also make use of fixed cells. These were prepared by washing twice with 37°C PBS, fixing in 4% paraformaldehyde for 10 minutes and again washing twice with 37°C PBS.

### 3. Imaging

We use an Andor Dragonfly 505 spinning disk confocal microscope. We use a 100X 1.49 NA CFI SR HP Apo TIRF oil immersion lens with a 1.5X magnification objective and a 60X 1.40 NA CFI apochromat Lambda oil immersion lens with a 2X magnification objective with a 1024 x 1024 pixel iXon 888 Life EMCCD camera. Using finite burst mode, we collect 1000 frames on each field of view for 40s with an exposure time of 10 ms. Illumination mode PD2 is used.

### 4. Image analysis

In the microscopy images, the granules appear as bright blobs against a darker background, as can be seen in Fig. 1B in the main text. Some G3BP-GFP remains in the cytoplasm as granules form, so the background is not completely dark. Due to the unreliability of transfection, the strength of the signal varies from cell to cell, which makes simple thresholding an unsuitable method of cell detection. Therefore, the Difference of Gaussian (DoG) method is used, which is better able to detect blobs against a complex background by searching for local maxima (6). This gives the coordinates of the centre-point of each granule in the image. We remove any granule with an area less than 100 pixels, as these are too small for our analysis. We estimate the area using a flood fill with a threshold of $0.5l_{max}$ where $l_{max}$ is the maximum intensity in the image.

The next step of the image analysis is edge detection. We express the boundary of a granule as the distance $D(\varphi)$ , from the centre-point to the edge as a function of angle $\varphi$ for a set of 400 angles evenly distributed between 0 and $2\pi$. The edge is the point of maximum directional gradient of the fluorescent signal $g = \nabla I \cdot \hat{r}$, where $I$ is the image intensity and $\hat{r}$ is the radial unit vector. The gradient at the pixel in row *i*, and column *j*, $\nabla I_{i,j}$, is calculated using a fourth order approximation in the x and y directions

$$\nabla I_{i,j} = \frac{1}{12}\left(I_{i-2,j} - 8I_{i-1,j} + 8\,I_{i+1,j} - I_{i+2,j},\ I_{i,j-2} - 8I_{i,j-1} + 8\,I_{i,j+1} - I_{i,j+2}\right). \tag{S11}$$

Once we have obtained $D(\varphi)$, we reject any boundary where there is a large discontinuous jump between two adjacent points. The main factor is due to overhangs where the granule cannot be represented by a radial function. This is often due to granule merging events. These rejections are evaluated on a frame-by-frame basis, giving a boundary pass rate for each granule. We remove a granule from further analysis if the boundary is rejected in more than 60% of the frames.

Once the edge is detected, a Fourier transform of $D(\varphi)$ gives the fluctuation spectrum with $\upsilon_q$ the magnitude of the $q_{th}$ fluctuation mode and $q$ a positive integer. These modes are visualised in Fig. 1F.

The approach described thus far assumes that the base-shape of the granule is spherical, but in our experiments we find the average granule shape is not always spherical in the timescale of the observation. To rectify this issue, we follow the approach by Pécréaux et al. (7) and expand our description of each fluctuation mode to include a fluctuating term $F_q$ and a constant term $C_q$, such that we can write

$$\upsilon_q(t) = F_q \cos\left(\omega_q t + \delta_q\right) + C_q. \quad \text{(S12)}$$

The fluctuating and constant components of the perturbations can be extracted from the measured fluctuations with

$$\left|F_q\right|^2 = \langle\left|\upsilon_q\right|^2\rangle - \left|\langle\upsilon_q\rangle\right|^2,$$

$$\left|C_q\right|^2 = \left|\langle\nu_q\rangle\right|^2. \quad \text{(S13)}$$

$\left|F_q\right|^2$ is then used in place of $\langle\left|\upsilon_q\right|^2\rangle$ as the fluctuation measure for calculating σ and κ.

We assume that the granules are imaged through the equatorial plane. Since the cells are quite flat, Z-stack confocal images of fixed cells show that the granules broadly occupy the same plane, justifying the assumption that the granules do not significantly diffuse into and out of the plane over the course of our measurements.

### 5. Spectrum fitting

The final step in the analysis is to find the values of σ and κ which give the best fit between the theoretical and measured spectra.

We have to be careful in the choice of how we define the best fit. An error function, $\varepsilon(\sigma, \kappa)$, gives a quantitative value for the quality of the fit. A poor choice of error function leads to an over-emphasis of certain parts of the spectrum. For example, a root mean squared approach, with

$$\varepsilon^2(\sigma, \kappa) = \sum_q \left(\left|F_{q,\,theo}\right|^2 - \left|F_{q,\,exp}\right|^2\right)^2, \quad \text{(S14)}$$

where $\left|F_{q,\,theo}\right|^2$ is the theoretical spectrum given by Eq. S10 and $\left|F_{q,\,exp}\right|^2$ the measured spectrum either with or without the correction given in Eq. S13, leads the minimiser to prioritise only the first orders (low $q$) and neglect the much smaller higher orders. This leads to a very poor overall fit. We improve this by introducing an error measure that is based on the error ratio between the values of theoretical spectrum $\left|F_{q,\,theo}\right|^2$ and the measured spectrum $\left|F_{q,\,exp}\right|^2$,

$$\varepsilon_{log} = \sum_q \left| log_{10}\left[\frac{\left|F_{q,\,theo}\right|^2}{\left|F_{q,\,exp}\right|^2}\right]\right|. \tag{S15}$$

We use a combination of parameter sweeps and L-BFGS-B algorithms (8) to minimise the fitting error, $\varepsilon_{log}$.

### 6. Simulations of rotating rigid bodies

We generate a simulated, quenched rigid body with a normalised surface deformation given by

$$u(\theta, \varphi, t) = \sum_{l=2}^{15} \sum_{m=-l}^{l} Y_{lm}(\theta, \varphi) U_{lm} e^{i\theta'} e^{i\varphi'}, \tag{S16}$$

where $U_{lm}$ is a random magnitude between 0 and 1 selected from the pink noise spectrum, and $(\theta, \varphi)$ is a random point on the unit sphere. The radius of the simulated droplet is taken to be 1. The choice of pink noise, instead of uniform (white) noise, ensures that the magnitude of the fluctuation decreases with $l$, yielding more physically plausible shapes when compared to stress granules. The rigid body then undergoes a series of 100 random rotations. Firstly, a random axis is generated, uniformly distributed on the unit hemisphere. Then a random rotation magnitude is selected from a normal distribution with a mean of 0 and a standard deviation of $2^{\circ}$. The equatorial cross-section is taken after each rotation in the series, and subject to the analysis described above, yielding an estimate of the spectrum that would be measured of a granule undergoing rotations. See Fig. 1F in the main paper.

1. S. A. Safran, Fluctuations of spherical microemulsions. *J. Chem. Phys.* **78**, 2073–2076 (1983).
2. W. Häckl, U. Seifert, E. Sackmann, Effects of Fully and Partially Solubilized Amphiphiles on Bilayer Bending Stiffness and Temperature Dependence of the Effective Tension of Giant Vesicles. *J. Phys. II* **7**, 1141–1157 (1997).
3. H. Engelhardt, H. P. Duwe, E. Sackmann, Bilayer bending elasticity measured by Fourier analysis of thermally excited surface undulations of flaccid vesicles. *J. Phys. Lett.* **46**, 395–400 (1985).
4. E. Becker, W. J. Hiller, T. A. Kowalewski, Experimental and theoretical investigation of large-amplitude oscillations of liquid droplets. *J. Fluid Mech.* **231**, 189–210 (1991).
5. N. Kedersha, *et al.*, G3BP–Caprin1–USP10 complexes mediate stress granule condensation and associate with 40S subunits. *J. Cell Biol.* **212** (2016).
6. D. Marr, E. Hildreth, S. Brenner, Theory of edge detection. *Proc. R. Soc. Lond. B Biol.*

*Sci.* **207**, 187–217 (1980).
7. J. Pécréaux, H.-G. Döbereiner, J. Prost, J.-F. Joanny, P. Bassereau, Refined contour analysis of giant unilamellar vesicles. *Eur. Phys. J. E* **13**, 277–290 (2004).
8. E. Ziegel, Numerical Recipes: The Art of Scientific Computing. *Technometrics* **29**, 501–502 (1987).
